# The spindle orienting machinery requires activation, not just localization

**DOI:** 10.1101/2022.06.29.498167

**Authors:** Kathryn E. Neville, Tara M. Finegan, Nicholas Lowe, Philip M. Bellomio, Daxiang Na, Dan T. Bergstralh

## Abstract

The orientation of the mitotic spindle at metaphase determines the placement of the daughter cells. Spindle orientation in animals typically relies on an evolutionarily conserved biological machine comprised of at least four proteins - called Partner of Inscuteable (Pins), Gαi, Mushroom body defective (Mud), and Dynein in flies - that exert a pulling force on astral microtubules and reels the spindle into alignment. The canonical model for spindle orientation holds that the direction of pulling is determined by asymmetric placement of this machinery at the cell cortex. In most cell types, this placement is thought to be mediated by Pins, and a substantial body of literature is therefore devoted to identifying polarized cues that govern localized cortical enrichment of Pins. In *Drosophila* neuroblasts, for example, this cue is thought to be Inscuteable, which helps recruit Pins to the apical cell surface. In this study we revisit the canonical model. We find that spindle orientation in the follicular epithelium requires not only Pins localization but also activation, which relies on direct interaction between Pins and the multifunctional protein Discs large. This mechanism is distinct from the one mediated by Inscuteable, which we find also has an activating step. Together our results show that the canonical model is incomplete. Local enrichment of Pins is not sufficient to determine spindle orientation; an activation step is also necessary.

## Introduction

The orientation of cell division is implicated in cell fate, as in the asymmetric division of neural progenitor cells, and in tissue architecture, as in the expansion of the *Drosophila* imaginal wing disc (reviewed in (Bergstralh et al., 2017)). Division orientation is generally determined by the orientation of the mitotic spindle at metaphase. Spindle orientation is governed by a suite of evolutionarily conserved factors comprised of Partner of Inscuteable (Pins; vertebrate LGN; nematode GPR1/2), Gαi, Mushroom body defective (Mud; vertebrate NuMA, nematode LIN-5), and Dynein.

A combination of genetic experiments and biochemical work has led to a generalized model in which Gαi acts as a membrane anchor for Pins, which in turn recruits Mud to the cortex. Direct binding between Pins/LGN and Mud/NuMA is mediated by tetratricopeptide repeats (TPRs) at the N-terminus of Pins/LGN (Du et al. 2001; Siller et al. 2006). Mud and Dynein together exert a pulling force on astral microtubules that reels the spindle into alignment. This model predicts that the key determinant of division orientation should be the placement of the pulling machinery at the cortex.

In most cell types, Pins/LGN is suggested to regulate the cortical position of the pulling machinery. For that reason, multiple studies have focused on the identification and characterization of cues that regulate Pins/LGN/GPR1/2 localization (reviewed in (Bergstralh et al., 2017)). In *Drosophila* neuroblasts, which are asymmetrically-dividing neural progenitor cells, this cue is thought to be a protein called Inscuteable (Kraut et al., 1996; Yu et al., 2000; Schaefer et al., 2000). In symmetrically-dividing epithelial cells, proposed Pins/LGN-positioning cues include the junctional proteins E-Cadherin and Afadin (in mammalian culture cells) and the multifunctional scaffolding protein Discs large (in flies, cultured mammalian cells, and chick neuroepithelium) (Gloerich et al., 2017; Carminati et al., 2016; Bergstralh et al., 2013a; Saadaoui et al., 2014; Chanet et al., 2017). All of these proteins are reported to interact with Pins directly. The proliferation of potential Pins-positioning proteins, particularly in epithelial cells, presents a problem: why are multiple factors necessary, especially in the same cell type?

The function of Pins is also somewhat enigmatic. Essentially, the model holds that it is an adaptor for an adaptor, linking Mud and therefore dynein to the cortex. The inclusion of a Pins-positioning factor like Discs large (Dlg) adds another adaptor to the mechanism. Again, there is a parsimony problem; why are so many adaptors necessary? This question is highlighted by literature showing that the number can be reduced: 1) During anaphase, NuMA anchors itself to the membrane through a cryptic motif that is uncovered by CDK1 phosphorylation (Seldin et al., 2013; Kiyomitsu and Cheeseman, 2013; Zheng et al., 2014; Kotak et al., 2013); and 2) the planar cell polarity protein Dishevelled can act as an alternative cortical anchor for Mud (Segalen et al., 2010).

These questions led us to revisit the canonical model and test its predictions, starting from the most fundamental. We find that the model is incomplete. Spindle orientation relies not only on the position of Pins but also on interaction with factors that promote its activity.

## Results

### Cortical Mud localization relies on Pins

Mitotic spindles in epithelial cells tend to orient along the plane of the tissue, meaning perpendicular to the apical-basal axis. In this way, divisions are oriented such that the two new daughter cells appear within the tissue (reviewed in (Bergstralh et al., 2013b)). The pulling machinery should therefore be expected to work at or adjacent to epithelial cell-cell borders (*i*.*e*. at the lateral cortex), and in agreement we have already shown that both Mud and Pins are lateral in mitotic follicle cells (Bergstralh et al., 2013a).

The canonical model holds that Pins acts as a cortical recruitment factor for Mud during mitosis (reviewed in (Morin and Bellaïche, 2011; Werts, 2011; Siller and Doe, 2009; Bergstralh et al., 2017; di Pietro et al., 2016)). Whether this holds true in *Drosophila* epithelia has not been tested outside of the pupal notum and larval wing disc, developmentally-related tissues in which Mud localizes to tricellular junctions in a Pins-independent manner (David et al., 2005; Bosveld et al., 2016; Bergstralh et al., 2016; Nakajima et al., 2019). We tested it in the *Drosophila* follicular epithelium (FE) and show here that cortical localization of Mud in mitotic follicle cells relies on Pins, whereas Mud localization at spindle poles is Pins-independent (Figure 1A). This pattern was previously observed in mitotic neuroblasts (Siller et al., 2006). Unexpectedly, Mud is also cortical in interphase follicle cells, and this localization relies on Pins (Figure 1B). Pins therefore has a general, rather than mitosis-specific, role in Mud localization in the follicular epithelium.

**Figure 1:**
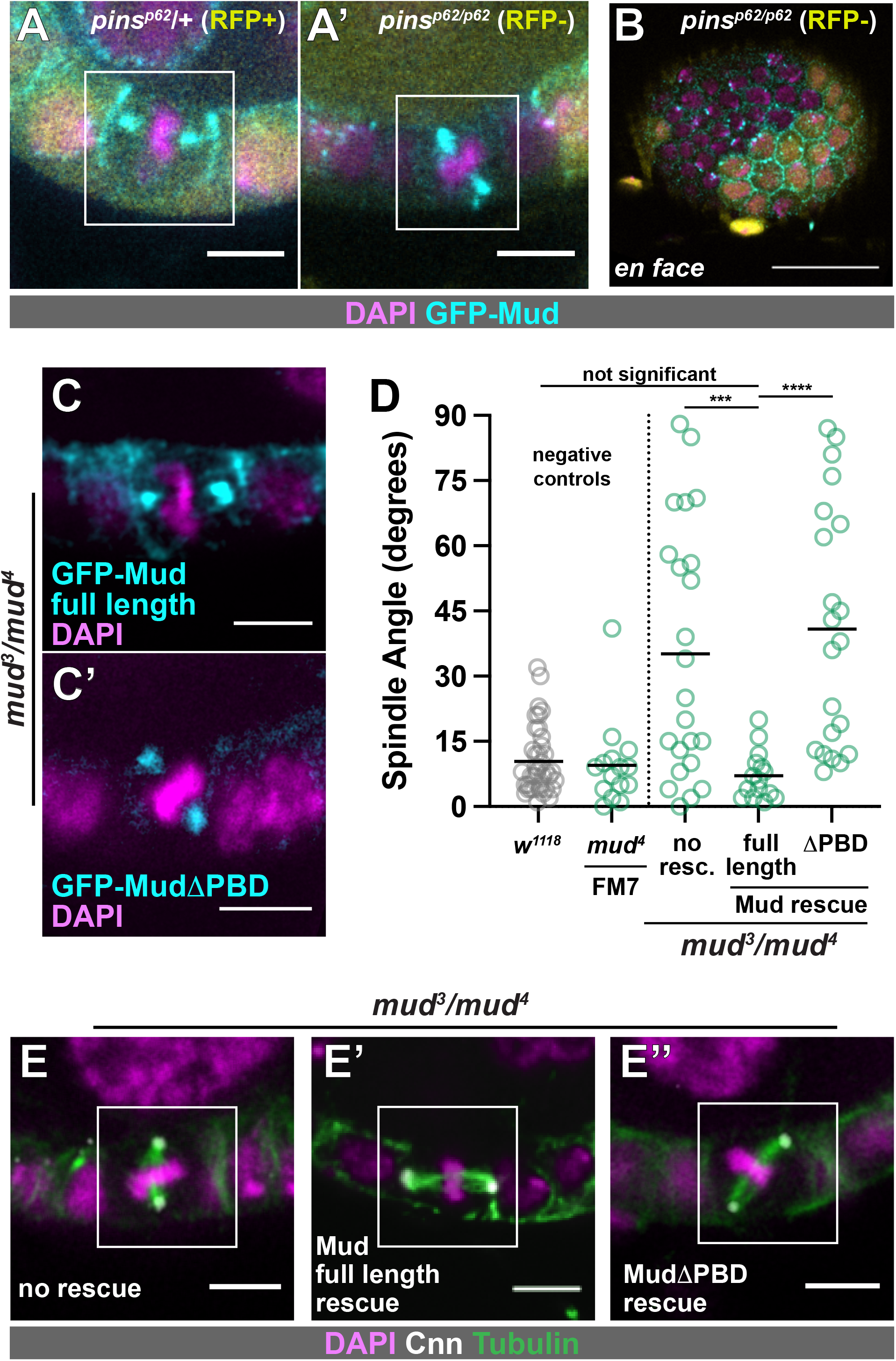
The cortical localization and spindle orienting function of Mud both rely on Pins. **A)** GFP:Mud is observed at spindle poles and at the lateral cell cortex in control mitotic follicle cells (A), but is only seen at the spindle poles in *pins*^*p62*^*/pins*^*p62*^ cells (A’). **B)** Interphase localization of Mud at the cortex is lost in *pins*^*p62*^*/pins*^*p62*^ follicle cells. **C, D, E)** Full length GFP:Mud shows the wild type pattern of localization in *mud*^*3*^*/mud*^*4*^ mitotic follicle cells (C) and rescues spindle orientation (D, E). GFP:Mud lacking the Pins binding domain localizes only at spindle poles (C’) in *mud*^*3*^*/mud*^*4*^ mitotic follicle cells and fails to rescue spindle orientation (D, E). Statistical significance in spindle orientation in this and subsequent figures was determined using the Mann-Whitney test. *\*\*\*\* p<0*.*0001, *** p<0*.*001*.

The Pins-binding domain (PBD) of Mud was previously mapped to a location between amino acids 1,928 and 1,982 (Izumi et al., 2006; Siller et al., 2006). We tested the importance of this region to spindle orientation in the FE using transgenes that encode the genomic promoter of Mud followed by either GFP-tagged full length Mud or Mud lacking the PBD (Bosveld et al., 2016). We expressed these transgenes in *mud*^*3*^*/mud*^*4*^ flies, in which endogenous Mud function is strongly impaired (Yu et al., 2006). In this background, full length GFP-Mud is observed at the cortex and at spindle poles and spindle orientation is rescued from random to the wild type distribution (Figure 1C-E). Contrastingly, GFP-MudΔPBD is observed at spindle poles but not the cortex, and fails to rescue spindle orientation (Figure 1C-E). One caveat to interpretation is that the PBD overlaps with a region of Mud that can bind microtubules (AA1,850–2,039), meaning that cortical localization may not be the only impairment caused by the deletion (Izumi et al., 2006; Siller et al., 2006). However, taken together our results are consistent with the canonical model; in the follicle epithelium Pins is required for the cortical localization and the spindle orientation function of Mud. A corollary to this finding is that the FE is a suitable system in which to interrogate the model.

### Pins may act as a cortical anchor for Mud, but this function is not obligate

We next characterized the subcellular localization of Pins in the FE. We generated transgenic flies expressing Pins:Tomato (Pins:Tom) under the control of the Pins genomic promoter. Pins:Tom is observed in all cell types in the egg chamber during the developmental stages at which follicle cells divide (prior to developmental Stage 7), and unlike Mud it can be detected past that point (Figure 2A). Pins:Tom overlaps with cortical actin; both are observed at the border between the FE and the interior germline cells and are more weakly detected at follicle cell-cell borders (Figure 2A). Unlike actin, however, Pins:Tom is not obvious at the basal cell surface (Figure 2B). We considered the possibility that Pins signal at the follicle cell-germline border is contributed by the germline cells, but clonal expression of Ubi-Pins-YFP shows that Pins localizes to the apical surface of follicle cells and accounts for the great majority of the signal at the border (Supplemental Figure 1A). In addition, although UAS-Pins-GFP is difficult to detect (a problem encountered by previous researchers) it is apically-enriched when expressed in the FE but not the germline (Supplemental Figure 1B) (Chanet et al., 2017). Together these results agree with previous work showing that the vertebrate ortholog of Pins (LGN) is apical in interphase epithelial cells, though we can also find Pins along lateral surfaces (Hao et al., 2010).

**Figure 2:**
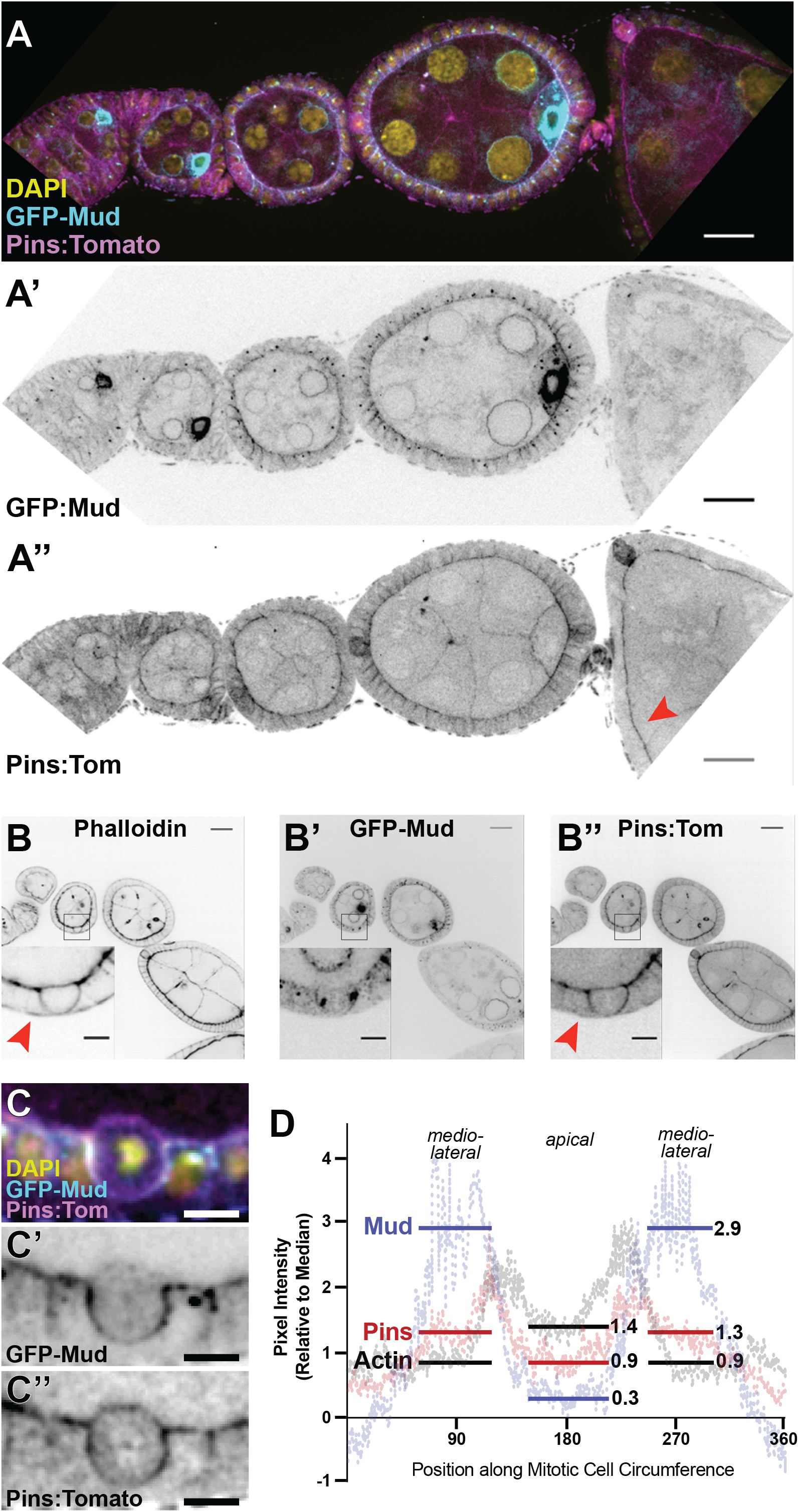
Expression and localization of Pins:Tom and GFP:Mud in interphase and mitotic follicle cells. **A)** Pins:Tom and GFP:Mud both appear enriched at the apical surface of follicle cells, and can be weakly detected at lateral surfaces. Enrichment of Mud at the oocyte nucleus has been reported previously (Zhao et al., 2012). Unlike GFP:Mud, Pins:Tom can be seen past Stage 6, the developmental point at which follicle cell division ceases. The red arrow points to a later (∼Stage 8) egg chamber. Scale bars = 20 microns. **B)** Pins and actin overlap extensively, though actin is observed at the basal surface (red arrow in B) whereas Pins is not (red arrow in B’’). Scale bars = 5 microns. **C**,**D)** Pins:Tom and GFP:Mud have overlapping but not identical localization patterns in mitotic follicle cells. Representative pictures are shown in (C). Quantification (9 cells) in (D). Scale bars = 5 microns.

The canonical model describes Pins as a recruitment factor for Mud, suggesting that the two proteins should localize together at the cortex. During interphase, GFP:Mud shows a similar pattern to Pins:Tom, though the signal is not strong enough to confirm colocalization (Figure 2A). Consistent with our prior work, we find that both proteins appear along the basolateral cortex in mitotic cells, with an enrichment just below the apical surface (we term this position “mediolateral”) and a decrease towards the basal surface (Figure 2B,C) (Bergstralh et al., 2013a). However, unlike Mud, Pins:Tom can also be distinguished at the apical cell surface (Figure 2C,D). Clonal expression of Ubi-Pins-YFP confirms that this signal originates primarily in the mitotic cell rather than the adjacent germline cell (Supplemental Figure 1C). Actin shows a similar pattern to Pins:Tom, but stronger enrichment at the apical surface (Figure 2C,D). A likely explanation is that this is at least in part because of cortical actin in the adjacent germline cell.

Together, these results show that in the follicular epithelium, the locations of Pins and Mud do not always correspond. This has also been shown in the wing disc and pupal notum, in which Pins is cortically-localized but does not control the location of Mud (Bosveld et al., 2016; David et al., 2005; Bergstralh et al., 2016). Our findings do not contradict a role for Pins in cortical anchoring of Mud. However, they suggest that this function is not obligate even in cells in which Mud relies on Pins for its cortical location.

### Relocalization of Pins at mitosis does not require a Pins-specific cue

We next addressed the question of how Pins relocalizes from the apical to the lateral cortex at mitosis. Previous work, including our own, has suggested that this an active change mediated by a cortical polarity cue (Hao et al., 2010; Bergstralh et al., 2013a). Another model to consider is that Pins relocalization occurs passively. During mitosis, the composition and material properties of the actin cortex undergo changes associated with cell rounding (Chugh and Paluch, 2018). The possibility that the change in Pins localization simply reflects these changes is suggested by the following observations: 1) Pins:Tom overlaps closely with actin throughout the cell cycle (Figure 2B and Supplemental Figure 2A); 2) both proteins are enriched at the mitotic cortex (Figure 2B); and 3) the unrelated transmembrane protein Basigin, a marker for follicle cell boundaries, is also cortically enriched at mitosis (Figure 2B and Supplemental Figure 2B) (Bergstralh et al., 2015).

We used UAS-Pins-myr-GFP, a myristoylated Pins variant developed by the Martin lab, to test between the active and passive models for Pins relocalization at mitosis (Chanet et al., 2017). We expressed Pins.myr-GFP in the FE and examined its localization and impact on spindle orientation. The active model predicts that the myristoyl modification might circumvent mechanisms that control asymmetric localization of Pins at the cortex, and that Pins-myr should therefore localize broadly around the plasma membrane. It does not. During interphase, Pins-myr is observed at the apical surface and lateral borders, though unlike Pins:Tom, Pins-myr signal extends evenly along the length of the epithelial cell-cell border rather than concentrating towards the apical side (Supplemental Figure 2A,C). During mitosis Pins-myr is enriched at the lateral cell-cell border and more weakly observed at the apical surface, a pattern very similar if not identical to Pins:Tom (Figure 3A,C). Myristoylated-RFP, a negative control, demonstrates similar localization to Pins:Tom and Pins-myr (Figure 3B,C and Supplemental Figure 2D). These results are consistent with the passive model. We also find that Pins-myr expression rescues spindle orientation in *pins* null tissue, indicating that it is able to substitute for wild-type Pins (Figure 3D-F). Together, our results indicate that the apparent relocalization of Pins from the apical to the lateral surface at mitosis – from which position it drives spindle orientation - can largely be explained as the placement of a membrane-anchored protein.

**Figure 3:**
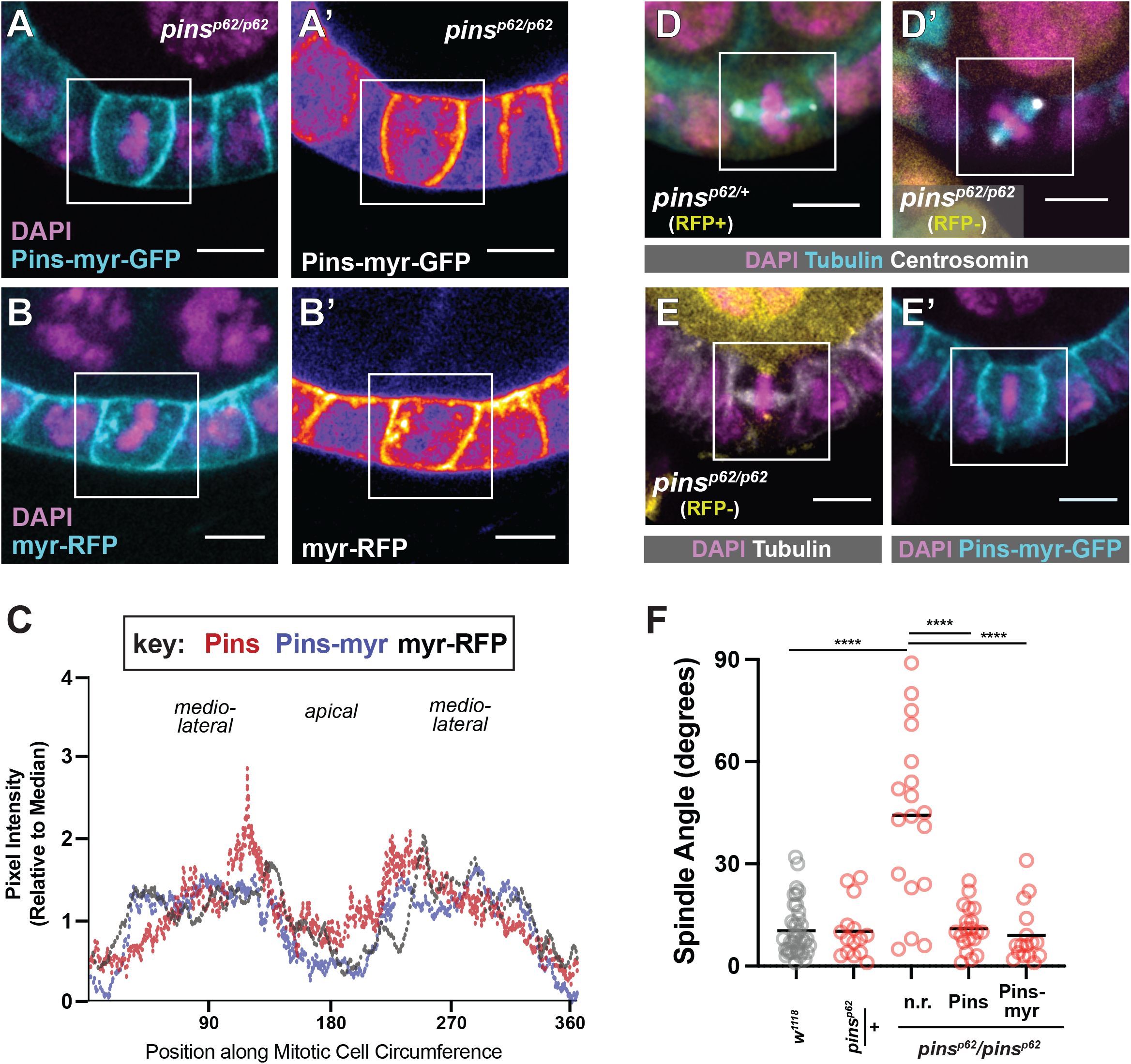
Pins does not require a Pins-specific cue to relocalize at mitosis. **A-C)** Pins-myr-GFP and myr-RFP show a similar localization pattern to Pins:Tom during mitosis. Representative pictures are shown in (A,B); a heatmap lookup table (A’,B’) emphasizes the relative enrichments of Pins-myr-GFP and myr-RFP along the cortex. Quantification (Pins-myr: 4 cells, myr-RFP: 2 cells) in (C). **(D-F)** Random spindle orientation in *pins*^*p62*^*/pins*^*p62*^ null mutant follicle cells is rescued by the expression of full-length Pins or Pins-myr-GFP. Representative pictures in (D,E) and quantification in (F). Scale bars = 5 microns. *\*\*\*\* p<0*.*0001*

### Pins is cortical in the absence of Dlg

The finding that membrane anchoring is sufficient to explain Pins localization at mitosis presents a challenge to our previous work, in which we suggested that relocalization relies on the epithelial polarity factor Discs large (Bergstralh et al., 2013a). Subsequent studies in the early *Drosophila* embryo, chick neuroepithelium, and unpolarized cultured cells indicated that Dlg is not just a cue but an anchor for Pins at the cell boundary (Saadaoui et al., 2014; Chanet et al., 2017). However, Pins/LGN is thought to be linked to the membrane by Gαi-GDP, raising the question of why a second anchor would be required (reviews (Morin & Bellaïche 2011; Werts 2011; Siller & Doe 2009; Bergstralh et al. 2017)). We therefore revisited the question of how Dlg participates in spindle orientation.

The spindle-orienting function of Dlg is thought to be mediated by direct interaction between the inactive C-terminal Guanylate Kinase (GUK) domain of Dlg and an evolutionarily conserved Pins phosphoserine (*Drosophila* S436, human S408) (Johnston et al., 2009; Zhu et al., 2011; Hao et al., 2010; Saadaoui et al., 2014). We confirmed this in the follicle epithelium in two ways: Firstly, we tested the role of Dlg. The *Dlg*^*1P20*^ allele encodes a truncated protein that is missing approximately one-third of its GUK domain and cannot bind to Pins *in vitro* (Bergstralh et al., 2016). We showed previously that spindles are misoriented in *Dlg*^*1P20*^ FE tissue, though apico-basal polarity is maintained (Bergstralh et al., 2013a). That analysis was performed using mitotic clones, raising the possibility that the spindle orientation defect could be caused by a second mutation on the *Dlg*^*1P20*^ chromosome. We tested this possibility using flies heterozygous for *Dlg*^*1P20*^ and *Dlg*^*2*^, a strong temperature-sensitive hypomorphic allele. We find that follicle cell spindles in *Dlg*^*1P20/2*^ egg chambers are misoriented at restrictive temperature (29C) (Figure 4A). These results confirm that disruption of the Dlg C-terminus causes spindle misorientation. Secondly, we examined the consequence of Pins phosphorylation. We generated transgenic flies encoding Pins, Pins-S436D (phosphomimetic), or Pins-S436A (unphosphorylatable) under UAS control and expressed these variants in *pins* null clones. We find that Pins and Pins-S436D both rescue spindle orientation, while Pins-S436A does not, in agreement with previous work (Figure 4B) (Johnston et al., 2009; Hao et al., 2010). These results are consistent with a model in which Dlg participates in spindle orientation through interaction with phosphorylated Pins.

**Figure 4:**
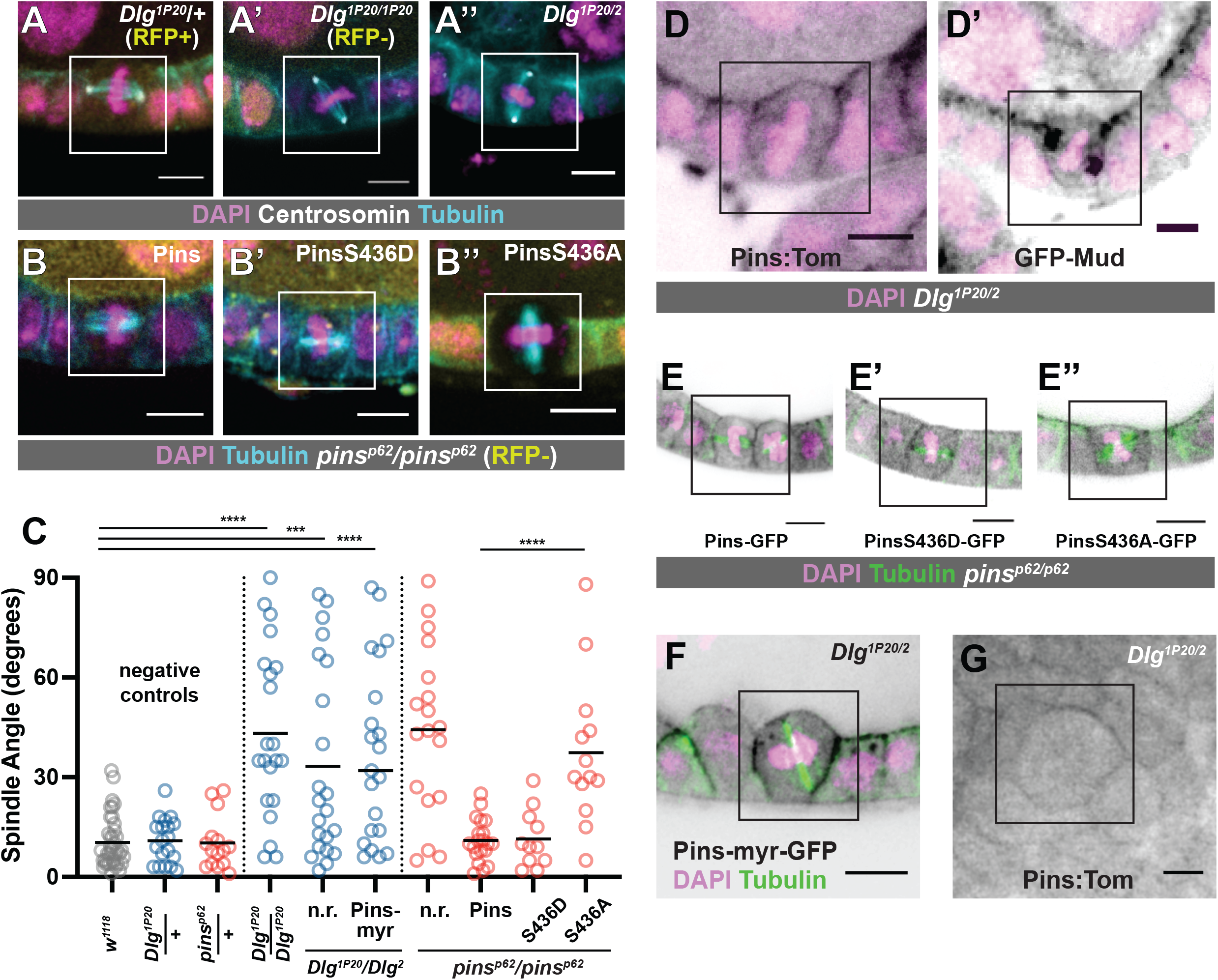
Dlg and Pins interact to promote spindle orientation in the follicle epithelium, but Dlg is not required for cortical localization of Pins. **A)** Spindle orientation is randomized in *Dlg*^*1P20*^*/Dlg*^*1P20*^ and *Dlg*^*1P20*^*/Dlg*^*2*^ mutant follicle cells. **B)** Random spindle orientation in *pins*^*p62*^*/pins*^*p62*^ null mutant follicle cells is rescued by the expression of full length Pins or phosphomimetic (S436D) Pins. Unphosphorylatable (S436A) Pins fails to rescue. **C)** Quantification of spindle orientation phenotypes. *\*\*\*\* p<0*.*0001, *** p<0*.*001*. **D)** Pins:Tom and GFP:Mud are cortical in *Dlg*^*1P20*^*/Dlg*^*2*^ mutant follicle cells. **E)** GFP-tagged Pins variants (control, phosphomimetic, unphosphorylatable) are observed at the mitotic cell cortex in *pins*^*p62*^*/pins*^*p62*^ null mutant follicle cells. **F)** Pins-myr-GFP fails to rescue spindle randomization in *Dlg*^*1P20*^*/Dlg*^*2*^ mutant follicle cells. Quantification in (C). **G)** Pins:Tom is cortical in an early embryo from a *Dlg*^*1P20*^*/Dlg*^*2*^ mother.

We next asked whether Dlg controlled Pins localization during mitosis. We showed previously that cortical recruitment of Pins in the FE is Dlg-independent, but a difficulty for interpretation is that these results relied on Pins-YFP, which is driven by a Ubiquitin promoter and therefore likely to be overexpressed (David et al., 2005; Bergstralh et al., 2013a). We repeated the experiment using Pins:Tom. We found that 1) Pins:Tom localizes to the mitotic cell cortex in *Dlg*^*14*^*/Dlg*^*14*^ mutant follicle cells, which are Dlg protein null, and in *Dlg*^*1P20*^*/Dlg*^*1P20*^ cells (Supplemental Figure 3A,B) and that 2) GFP-Mud and Pins:Tom localize to the mitotic cortex in *Dlg*^*1P20*^/*Dlg*^*2*^ mutant follicle cells (Figure 4D). Additionally, Pins, Pins-S436D, and Pins-S436A are all observed at the mitotic cortex in *pins* null clones (Figure 4E). These results indicate that the spindle orienting function of Dlg is not through cortical anchoring of Pins. To test this even further, we expressed Pins-myr in *Dlg*^*1P20*^/*Dlg*^*2*^ mutant follicle cells. While the localization of this variant is identical to Pins-myr in a control background, it fails to rescue spindle orientation (Figure 4C,F). Together, our findings indicate that while Dlg is important for spindle orientation in the FE, the effect is not through redistribution of Pins.

We also examined the relationship between Pins and Dlg in the *Drosophila* embryo. A previous study showed that cortical localization of Pins in embryonic epithelial cells is lost when *Dlg* mRNA is knocked down (Chanet et al., 2017). We find that Pins:Tom localizes to the cortex in early embryos from *Dlg*^*1P20*^/*Dlg*^*2*^ mothers, indicating that Pins localization in the embryo does not require direct interaction with Dlg. We therefore speculate that Dlg plays an additional role in that tissue, upstream of Pins (Figure 4G).

### Khc-73 is not required for spindle orientation in the follicular epithelium

Another proposed explanation for the role of Dlg in spindle orientation is that it facilitates interaction between Pins and a microtubule-capturing activity. This has been studied an artificially-polarized S2 (cultured) cell system, in which the microtubule-capturing factor is the plus-end directed motor Khc-73 (Siegrist and Doe, 2005; Johnston et al., 2009; Lu and Prehoda, 2013). Khc73 is also part of a Pins/Dlg pathway that rescues division orientation in Inscuteable-mutant neuroblasts (Siegrist and Doe, 2005). Given that epithelial cells express Dlg but not Inscuteable, the possibility that this rescue pathway is the default in epithelia is particularly attractive. We tested it in follicle cells using 1) flies transheterozygous for the null alleles *Khc-73*^*149*^ and *Khc-73*^*193*^, 2) expression of Khc73 shRNA in the follicular epithelium; and 3) mitotic clones mutant for the null allele *Khc73*^*3-3*^ (Liao et al., 2018; Zajac and Horne-Badovinac, 2022). Spindle angles were normal in all of these conditions (Figure 5A,C).

**Figure 5:**
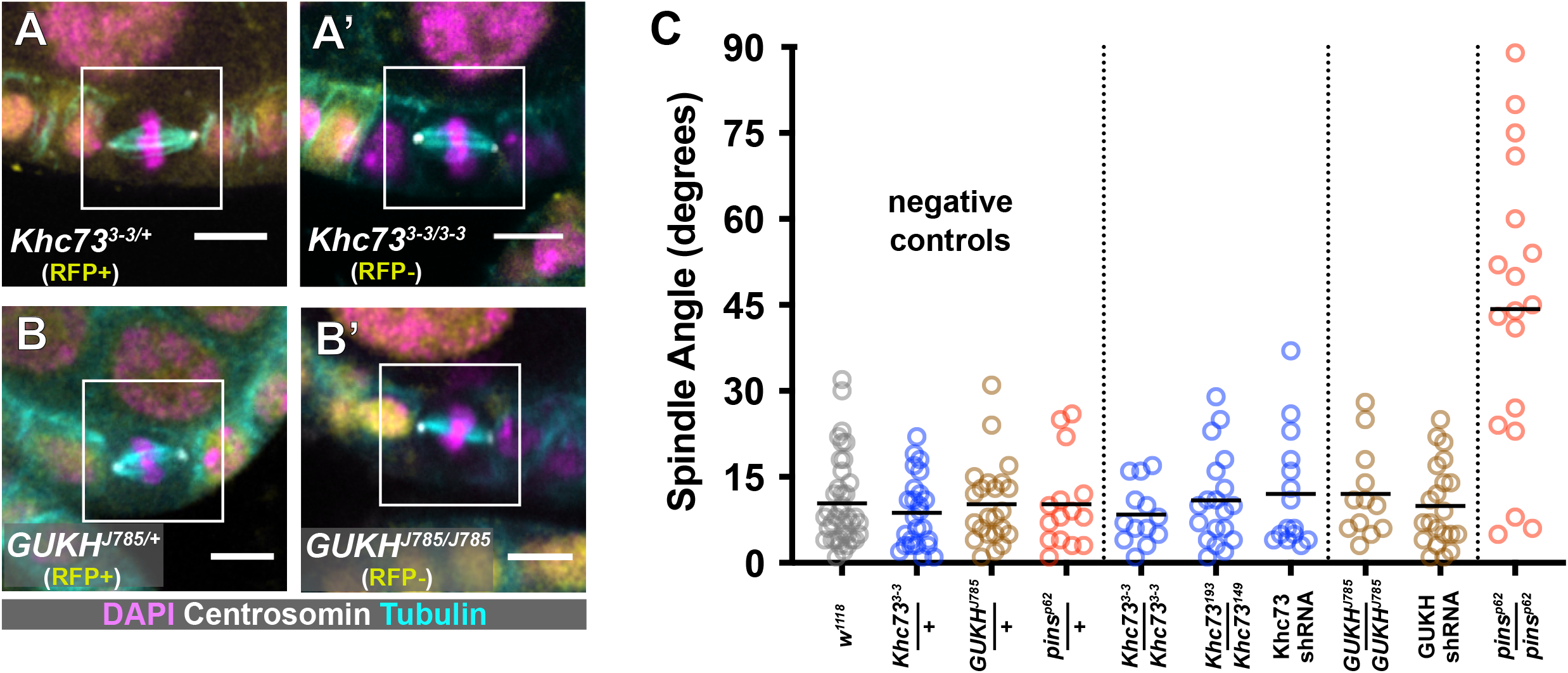
Putative microtubule-capturing factors are not required for spindle orientation in the follicular epithelium. **A-C)** Multiple genetic approaches were used to test a role for either Khc73 or GUKHolder in metaphase spindle orientation. Representative pictures are shown in (A) and (B). Quantification in (C). Spindle angle randomization in *pins*^*p62*^*/pins*^*p62*^ null mutant follicle cells is included as a positive control for comparison.

Dlg is also proposed to link the spindle orienting machinery to another microtubule capturing protein called GUKHolder (GUKH), though loss of GUKH function has a negligible effect on spindle orientation in neuroblasts (Golub et al., 2017). We examined spindle orientation in follicle cells mutant for *GUKH*^*J785*^, which encodes a premature stop at Q520. (Full length isoforms are >1700AA). This allele causes failed projection of photoreceptor axons and lethality (Berger et al., 2008). Spindle orientation was normal in *GUKH*^*J785*^ cells and likewise unaffected by expression of shRNA targeting *GUKH* transcript, though we do not have direct evidence of knockdown (Figure 5B,C). Together, these findings show that Khc73 is dispensable for spindle orientation in the follicle epithelium and do not support a role for GUKHolder.

### Two patterns of Inscuteable localization predict spindle orientation

Our findings indicate that Discs large promotes Pins-dependent spindle orientation without acting as a localization factor or a link to microtubule-capturing molecules. They also show that, in contradiction to the model, Pins localization is not sufficient to determine the activity of the pulling force. Another way to test this is to move Pins to a different location on the cortex.

A potential strategy for relocalizing the spindle machinery is ectopic expression of Inscuteable, which is required for apical localization of Pins in neuroblasts (Yu et al., 2000; Schaefer et al., 2000). This manipulation has a well-established impact on the position of the pulling force; ectopic expression of Inscuteable in any of three *Drosophila* epithelia - the embryonic ectoderm, larval neuroepithelium, and follicular epithelium - causes reorientation of mitotic spindles such that they are parallel, rather than perpendicular, to the apical-basal axis (Kraut et al., 1996; Bowman et al., 2006; Egger et al., 2007; Bergstralh et al., 2015). Our own earlier investigation of UAS-Inscuteable expression in the FE was performed using the mosaic GAL4 drivers GR1-GAL4 and T155-GAL4 and relied on visualization of Inscuteable (by immunostaining) at the apical cell cortex to confirm expression (Bergstralh et al., 2015). We repeated the experiment using Traffic jam-GAL4, which is less mosaic across a given egg chamber, though in our hands generally weaker than other drivers. In agreement with our previous work, cells with a strong (∼4-fold > median) apical enrichment of Inscuteable (Insc^A^) demonstrate perpendicular spindle orientation (Figure 6A,B,F and Supplemental Figure 1A) (Bergstralh et al., 2015). However, we also identified a population of cells that demonstrates both lateral and apical localization of Inscuteable (Insc^B^) (Figure 6C,D). In an exceptional instance, it is only lateral (Supplemental Figure 1B). Spindle orientation is normal in this group (Figure 6F). We considered whether Insc^A^ cells resolve to become Insc^B^, but consider this possibility unlikely because we observed misoriented divisions in this study and previous work (not shown and (Bergstralh et al., 2015)).

**Figure 6:**
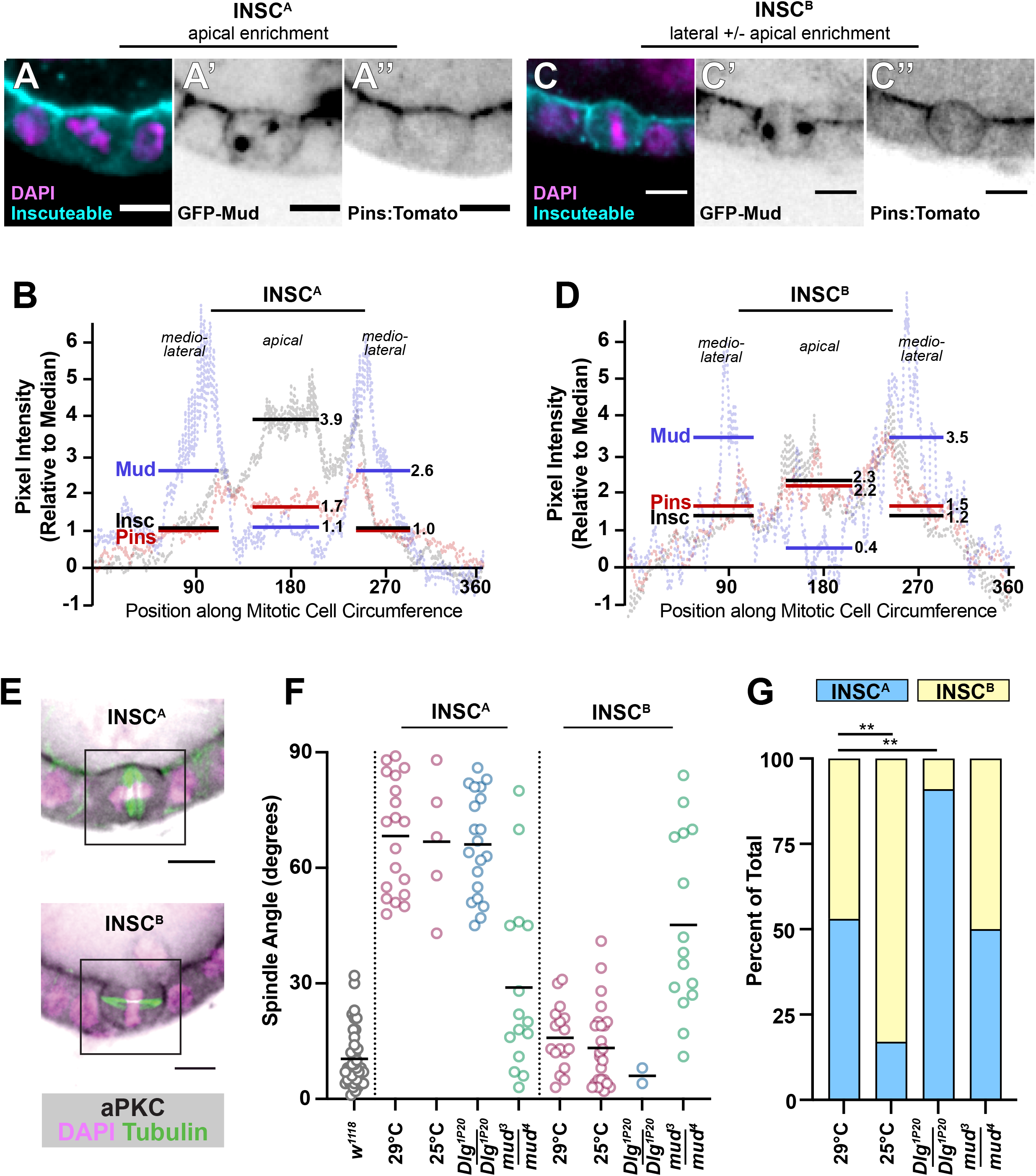
Inscuteable and Dlg function as distinct activators of the pulling machinery. **A-D)** Two distinct patterns of Inscuteable localization are observed in the follicular epithelium when UAS-Inscuteable is driven by Traffic jam-GAL4. (A,C) In the first pattern (Insc^A^), Inscuteable is highly enriched at the apical cell surface. (B,D) In the second pattern (Insc^B^), Inscuteable is observed at both the lateral and apical cortex. Mud and Pins localizations are also shown. Representative pictures are shown in (A,B) and quantifications (Insc^A^: average of 5 cells, Insc^B^: average of 4 cells) in (C,D). The example shown in (A) is an extreme case and is used to highlight the finding that Pins and Mud localizations do not strictly overlap. **E)** aPKC localization in Insc^A^ and Insc^B^ follicle cells. **F)** Quantification of spindle orientation phenotypes shows that mitotic spindles are reoriented in Insc^A^ cells, but not Insc^B^ cells. Reorientation relies on Mud but not on the interaction between Pins and Dlg. **G)** The ratio of Insc^A^ to Insc^B^ is impacted by temperature and by the interaction between Pins and Dlg. Significance was determined using the Chi-square test. *\*\* p<0*.*01*

We speculate that the two populations reflect different expression levels, meaning that there is a threshold of expression over which Inscuteable causes reorientation. Two lines of evidence support this possibility. Firstly, ectopic expression of Inscuteable in the embryonic ectoderm causes a reliable reorientation of spindle angles rather than a bimodal distribution, consistent with the possibility that the threshold is met in that tissue (Kraut et al., 1996; Bergstralh et al., 2015). Secondly, we found that spindle reorientation was less common when UAS-Inscuteable flies were maintained at 25°C rather than 29°C (Figure 6E,F). The UAS-GAL4 system is more efficient at the higher temperature.

### The pulling force must be activated, not just localized

Although Inscuteable can reorient epithelial cell spindles, a careful examination of the spindle orienting machinery in epithelial cells expressing Inscuteable has not been undertaken. We first tested the influence of Inscuteable on cortical polarity. In neuroblasts, Inscuteable and the Par complex proteins aPKC and Bazooka rely on one another for apical localization (Wodarz et al., 1999; 2000; Yu et al., 2000; Schaefer et al., 2000). In wild type follicle cells aPKC is an apical polarity marker at interphase but loses its polarized enrichment during mitosis (Bergstralh et al., 2013a; Morais-de-Sá and Sunkel, 2013). In agreement with interdependence between Inscuteable and the Par complex, we find that aPKC is stabilized at the apical cortex in Insc^A^ cells but enriched at the lateral cortex in Insc^B^ cells (Figure 6E). This finding is consistent with an Inscuteable-expression threshold model; below the threshold, Pins dictates lateral localization of Inscuteable and aPKC. Above the threshold, Inscuteable dictates apical localization of Pins and aPKC.

The canonical model predicts that Pins and Mud should be recruited to the apical surface in Insc^A^ cells, from which position they would draw the spindle into alignment along the apical-basal axis, and that Pins and Mud should be primarily lateral in Insc^B^ cells, where they would promote wild type spindle orientation (perpendicular to the apical-basal axis). These predictions are not met. Instead, we find that Pins is apically enriched in both Insc^A^ and Insc^B^ cells, and this enrichment is roughly equivalent in both populations (Figure 6A-D). Mud looks markedly different to Pins. Although a weak enrichment is observed at the apical surface in Insc^A^ cells, Mud is most prominent at the mediolateral regions in both Insc^A^ and Insc^B^ cells (Figure 6A-D).

These results conflict with the canonical model. If spindle orientation is predicted only by the location of Pins, Insc^A^ and Insc^B^ cells should both be expected to orient their spindles along the apical-basal axis, as in neuroblasts. If spindle orientation is predicted only by the location of Mud, Insc^A^ and Insc^B^ cells should both be expected to orient their spindles along the tissue plane, as in the wild type. Our results show that the strongest predictor of spindle orientation in these cells is neither Mud nor Pins, but rather Inscuteable; if it is highly enriched at the apical surface (Insc^A^) spindles are reoriented, but if it is not (Insc^B^) then spindle angles are normal.

Since Mud localization is not an accurate predictor of spindle orientation, whereas Inscuteable is, a possibility to consider is that Inscuteable exerts a Mud-independent effect. However, we find that spindle orientation is randomized in both both Insc^A^ and Insc^B^ cells when Mud is genetically removed (Figure 6F). This finding indicates that in Insc^A^ cells Inscuteable impacts both the location and the activity of Mud-dependent pulling.

We have already shown that activation of pulling in the FE requires Dlg, and an obvious possibility is therefore that Inscuteable promotes spindle reorientation in Insc^A^ cells by changing the location of Dlg. We have been unable to test this directly because Dlg is expressed in germline cells as well as the FE, meaning that it can appear to be apical even in wild type tissue. However, to find out whether Dlg is required for Inscuteable-mediate spindle orientation we expressed Inscuteable in *Dlg*^*1P20*^/*Dlg*^*1P20*^ mutant tissue. Not only does Inscuteable reorient spindles in this tissue, but the proportion of Insc^A^ cells is significantly higher (Figure 6F,G and Supplemental Figure 4C). In other words, Dlg acts to shift the threshold towards the Insc^B^ pattern. We interpret this to mean that interaction between Pins and Dlg, which is required for pulling, stabilizes the lateral pulling machinery even if Dlg is not a direct anchor.

Taken together with the results described above, these findings show that Inscuteable and Dlg mediate distinct and competitive mechanisms for activation of the spindle-orienting machinery in follicle cells.

## Discussion

### Limitations of this Study

A technical challenge to undertaking this study is that current fluorescent molecules - including those introduced here - are exceedingly weak and liable to photobleaching. (Pins-myr-GFP is an exception). This issue presents a barrier to live imaging, meaning that temporal resolution is unavailable. We are also unable to convincingly determine whether cortical Pins is decreased (as opposed to simply absent), when it is unable to interact with Dlg; a decrease is predicted by work in chick embryo neuroepithelial cells showing that a C-terminal fragment of LGN localization is ∼33% less cortical in the unphosphorylatable condition (Saadaoui et al., 2014). However, we find that even when Pins is anchored to the membrane by myristoylation it fails to rescue spindle orientation in *Dlg*^*1P20*^/*Dlg*^*2*^ mutant tissue, meaning that a reduction of Pins is unlikely to be the only effect responsible for randomization.

### Additional Function for Pins?

While it is not a focus for our study, our work provides support for the possibility that Pins has an additional function outside of spindle orientation. This is already suggested by the fact that strong mutants of *pins* are lethal to the organism, whereas strong mutants of *mud* are viable (Yu et al., 2000; Schaefer et al., 2000; Yu et al., 2006). Additionally, in the developing mouse epidermis, the vertebrate ortholog of Pins (LGN) acts to promote stratification by facilitating daughter cell placement subsequent to metaphase, perhaps implicating LGN in the regulation of cell-cell adhesion (Lough et al., 2019). We show here that whereas Mud expression in the follicular epithelium ends at developmental Stage 6, coinciding with the end of follicle cell divisions, Pins continues to be detected past that point (Bergstralh et al., 2013a). This suggests that Pins function is not limited to a role in proliferation. In addition, we find that while the LGN S408D (*Drosophila* 436D) variant is reported to act as a phosphomimetic, expression of this variant has no obvious effect on division orientation (Johnston et al., 2012). What then is the purpose of this modification? One possibility to consider is that the phosphorylation regulates the spindle orientation role of Pins, whereas unphosphorylated Pins plays a different role.

### Dlg and Inscuteable function as distinct activators for the pulling machinery

Previous work, including our own, has concentrated on identifying mechanisms that regulate the position of the spindle-orienting machinery. Work presented here shows that localization of either Pins or Mud is not always sufficient to predict pulling. Firstly, while Pins is required for cortical localization of Mud in the follicular epithelium, Pins and Mud localizations do not strictly overlap, meaning that Pins is not an obligate Mud anchor. We see evidence for this in wild type mitotic follicle cells, in which Pins appears slightly apical whereas Mud does not, and in follicle cells expressing Inscuteable, in which the pattern of Pins and Mud localizations are markedly different. Because cortical localization of Mud/NuMA has proven difficult to visualize, Pins location has been used as a proxy for the position of the entire pulling machinery (Dimitracopoulos et al., 2020). While this may very well be the case in some cell types, our work suggests that it is not a rule that can be automatically applied. Secondly, we find that even Mud/NuMA localization is not necessarily a reliable predictor for spindle orientation (Figure 6). Although elegant optogenetics work shows that cortical anchoring of NuMa is sufficient to direct pulling in unpolarized HeLa cells, we find that in the follicular epithelium at least Mud activity must also be regulated by additional factors (Okumura et al.).

One of these factors is Discs large. We tested two models for the function of Dlg in Pins-mediated spindle orientation in the follicular epithelium. The first model, based in part on our own previous work, is that Dlg acts as a cortical anchor for Pins, and is therefore important for relocalization of Pins from the apical to the lateral cortex at mitosis (Bergstralh et al., 2013a). We now show that this model is incorrect, partly because Pins relocalization at mitosis does not require a Pins-specific mechanism, but only a membrane anchor. Gαi may serve that purpose. The second model is that Dlg links Pins with Khc73 or GUKHolder. We find that neither molecule is important for spindle orientation in the FE. Our findings indicate that instead of a localization cue or a link to microtubule-capturing factors, Dlg acts as an activator for Pins-mediated pulling in the follicular epithelium. This mechanism is distinct from and can compete with Inscuteable, which we show here is not only a cortical localization factor for the spindle-orienting machinery but also an activator.

The two mechanisms are not likely to work in the same way. Dlg binds to a conserved phosphosite in the central linker domain of Pins and should therefore allow for Pins to simultaneously interact with Mud (Johnston et al., 2009). Contrastingly, binding between Pins and Inscuteable is mediated by the TPR domains of Pins, meaning that Mud is excluded (Culurgioni et al., 2011; 2018). While a stable Pins-Inscuteable complex has been suggested to promote localization of a separate Pins-Mud-dynein complex, our work raises the possibility that it might also or instead promote activation.

The finding that there are two distinct mechanisms for activating pulling raises the question of how and for what purpose they evolved. Phylogenetic analysis indicates that the interaction between Pins and Dlg is evolutionarily ancient; it is predicted to have evolved in Cnidaria, predating the appearance of Inscuteable orthologs (Schiller and Bergstralh, 2021). It is therefore tempting to speculate that Inscuteable-mediated spindle orientation evolved to promote cell type diversity, particularly in the more complex bilaterian nervous system.

Together, our results show that the canonical model that explains the mitotic spindle orientation is incomplete. Spindle orientation relies not only on the localization, but also the activation of the pulling machinery.

## Figure Legends

**Supplemental Figure 1 – Related to Fig 2:**
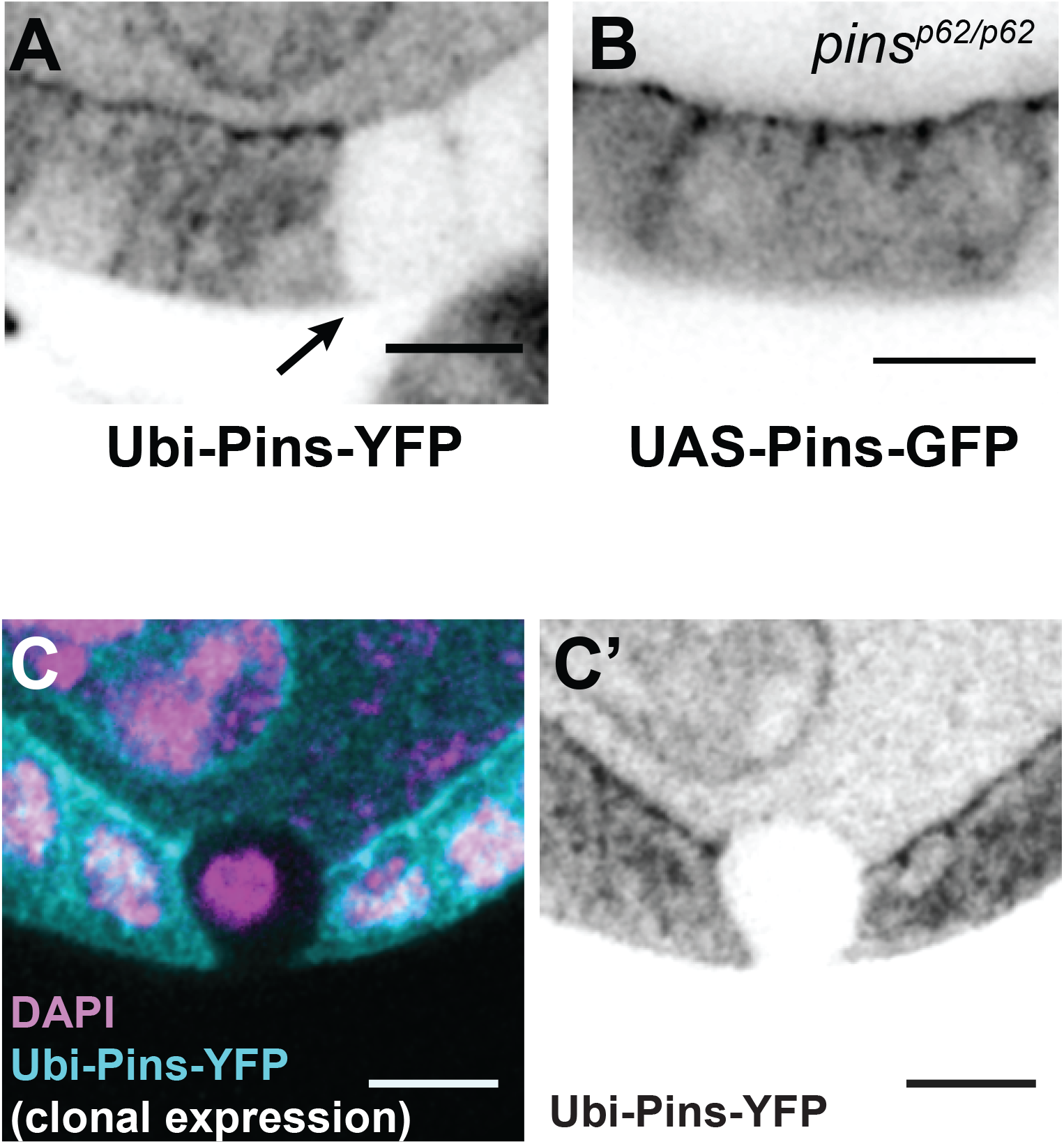
Pins signal at the border between interior germline and exterior follicle cells reflects apical localization in follicle cells. **A)** Clonal expression of Ubi-Pins-YFP shows that most signal at the border between germline and follicle cells originates in the follicle epithelium. The arrow marks the clone border. **B)** Pins-GFP appears at the apical surface when expressed only in the follicle epithelium. **C)** Clonal expression of Ubi-Pins-YFP shows that signal at the border between the germline and a mitotic follicle cell originates primarily in the mitotic cell. In this field of view, only the mitotic cell lacks expression of Ubi-Pins-YFP.

**Supplemental Figure 2 – Related to Fig 3:**
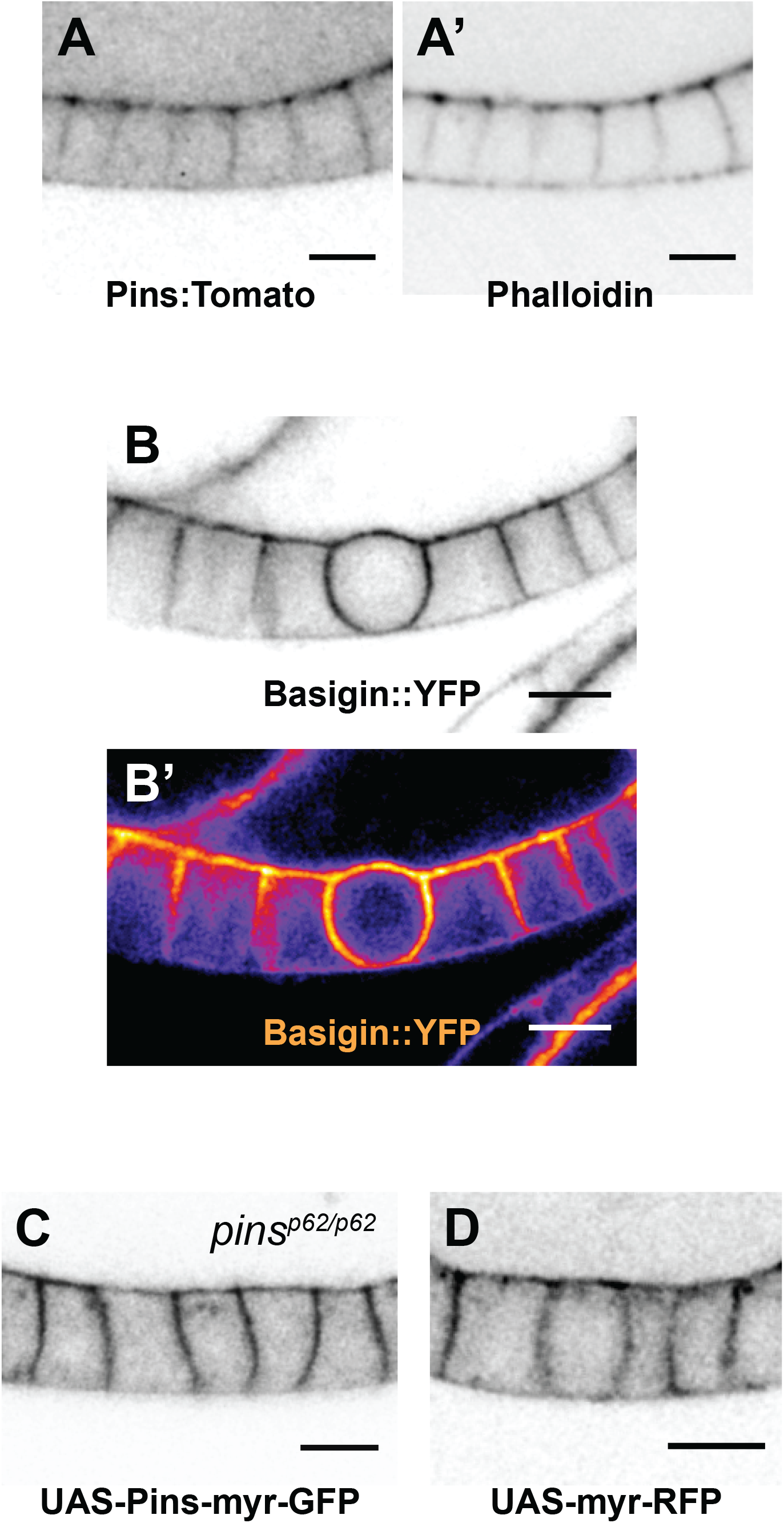
Localization of cortical proteins in the follicular epithelium **A)** Pins:Tom and actin overlap at the apical and lateral cortex during interphase. **B)** Basigin::YFP is enriched in a mitotic cell. Both a b/w and heat map image are shown to highlight enrichment. The protein can be found at the borders of germline and follicle cells. **C**,**D**) UAS-myr-Pins-GFP (B) and UAS-myr-RFP localization (C) show similar localization in interphase follicle cells.

**Supplemental Figure 3 – Related to Fig 4:**
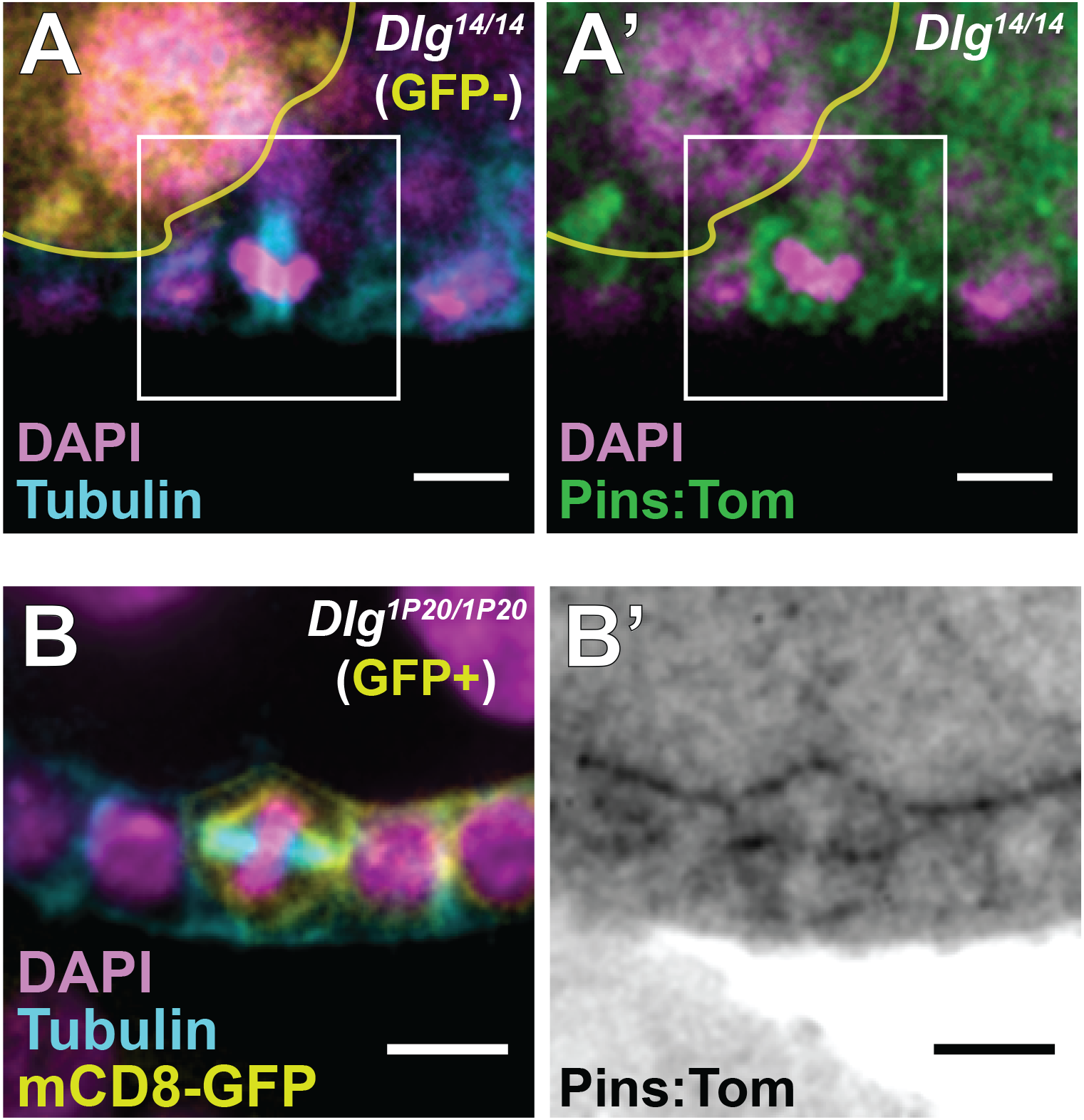
Pins:Tom is observed at the cell cortex in **A)** a mitotic *Dlg*^*14*^*/Dlg*^*14*^ cell and **B)** a mitotic *Dlg*^*1P20*^*/Dlg*^*1P20*^ cell. For the former, mutant tissue is marked by the absence of RFP. For the latter, mutant tissue is marked by the presence of mCD8-GFP.

**Supplemental Figure 4 – Related to Fig 6:**
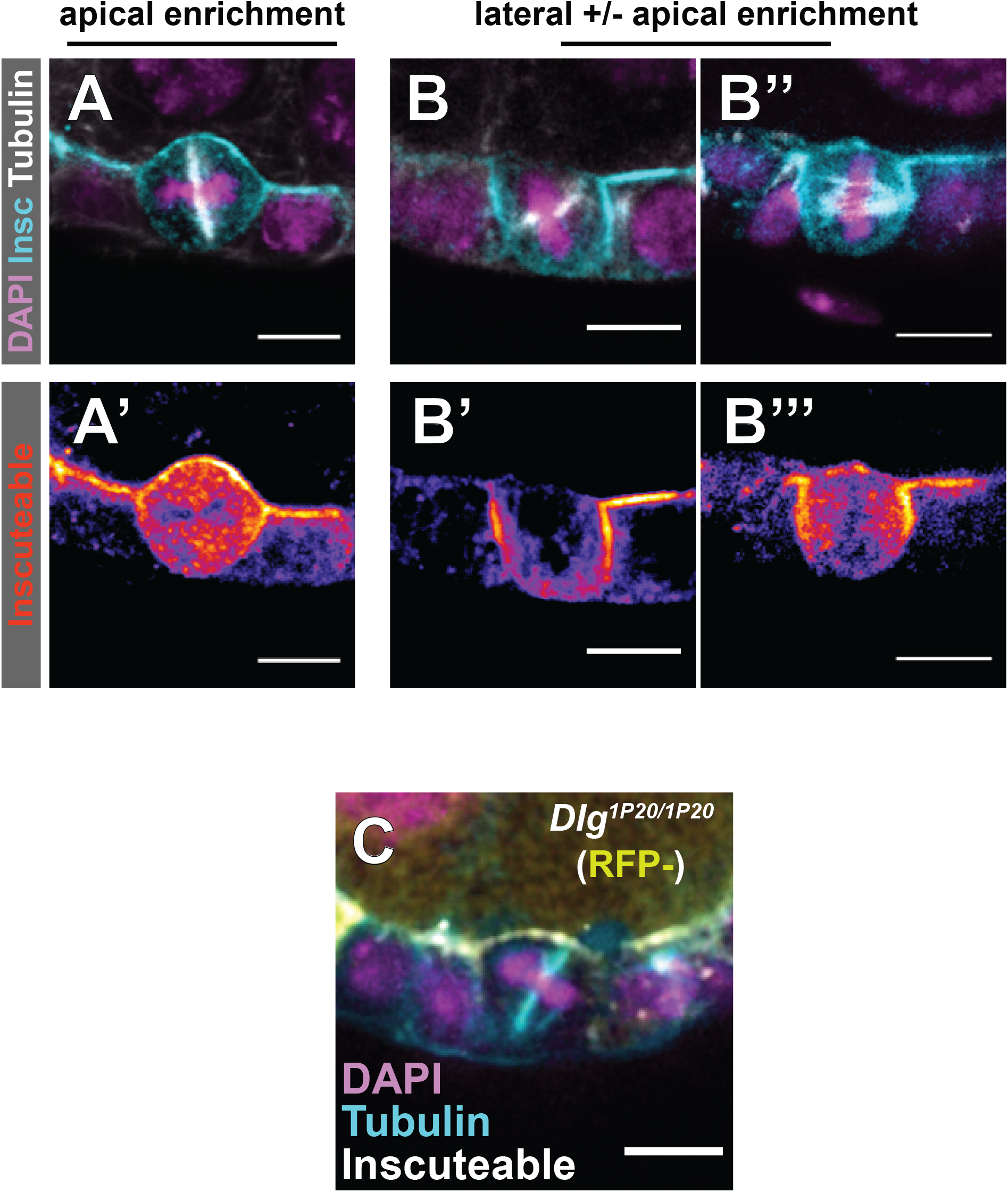
Insc^A^ and Insc^B^ cells are distinguished by the pattern of Inscuteable localization. **A)** In Insc^A^ cells, Inscuteable is strongly enriched at the apical surface. **B)** Insc^B^ cells show a stronger enrichment of Inscuteable at lateral surfaces. In an extreme example, Inscuteable is only lateral. **C)** Mitotic spindle reorientation in a Dlg^1P20^/Dlg^1P20^ cell expressing Inscuteable. Quantification in Figure 6F.

## Acknowledgments

We are grateful to the Rochester-Manchester-Carolina Meeting Group, the Rochester Invertebrate Group, Holly Lovegrove, and to Bergstralh lab members for comments. Rebecca (Becky) Bastock suggested clonal expression of Ubi-Pins-YFP. We thank Allison Zajac (Horne-Badovinac lab), Adam Martin, Pejmun Haghighi for sharing flies.

## Author Contributions

DTB conceived the project. DTB designed the experiments. DTB, KEN, and TMF performed the experiments. NL, PB, and DN generated transgenic flies. DTB wrote the manuscript.

## Materials and Methods

### *Drosophila* genetics

A list of alleles and transgenes used in this study is found in Table 1. We thank the Transgenic RNAi Project at Harvard Medical School (NIH/NIGMS R01-GM084947) for providing shRNA lines. Ectopic protein expression was accomplished using the UAS-GAL4 system (Brand and Perrimon, 1993). Expression was driven by Traffic Jam-GAL4 (Olivieri et al., 2010).

**Table 1.** *Drosophila* genetics.

| **Mutant Allele or Transgene** | **Citation** | **Stock Number or Lab of Origin** |
| --- | --- | --- |
| *mud*<sup>3</sup> | (J. X. Yu, Guan, & Nash, 2006) | Nash lab |
| *mud*<sup>4</sup> | (J. X. Yu et al., 2006) | Nash lab |
| *dlg*<sup>1P20</sup> aka *dlg*<sup>18</sup> | (Woods & Bryant, 1989) | Bloomington 36279 |
| *dlg*<sup>14</sup> | (Woods & Bryant, 1989) | Bloomington 36283 |
| *dlg*<sup>2</sup> | (Perrimon, 1988) | Bloomington 36278 |
| *pins*<sup>p62</sup> | (F. Yu, Morin, Cai, Yang, & Chia, 2000) | Chia lab |
| *GUKH*<sup>J785</sup> | (Berger et al., 2008) | Kyoto 109913 |
| *Khc73*<sup>3-3</sup> | (Zajac & Horne-Badovinac, 2022) | Horne-Badovinac lab |
| *Khc73*<sup>193</sup> | (Liao et al., 2018) | Haghighi lab |
| *Khc73*<sup>149</sup> | (Liao et al., 2018) | Haghighi lab |
| UAS-GUKH-shRNA TRiP.HMC03681 | (Perkins, Holderbaum, Tao, Hu, & Sopko, 2015) | Bloomington 55858 |
| UAS-Khc73-shRNA TRiP.HMS01624 | (Perkins et al., 2015) | Bloomington 367333 |
| UAS-Pins-GFP | this study |  |
| UAS-PinsS436D-GFP | this study |  |
| UAS-PinsS436A-GFP | this study |  |
| UAS-Inscuteable | (Kraut, Chia, Jan, Jan, & Knoblich, 1996) | Knoblich lab |
| UAS-Pins-myr-GFP | (Chanet, Sharan, Khan, & Martin, 2017) | Martin lab |
| UAS-myr-RFP | donated to the Bloomington Stock Center by Henry Chang | Bloomington 7118 |
| Ubi-Pins-YFP | (David et al., 2005) | Bellaïche lab |
| GFP-Mud | (Bosveld et al., 2016) | Bellaïche lab |
| GFP-MudΔPBD | (Bosveld et al., 2016) | Bellaïche lab |
| Pins:Tomato | this study |  |
| Basigin::YFP | (Lovegrove, Bergstralh, & St Johnston, 2019) | Kyoto 115365 |

### Mitotic clones

Clones were induced by incubating larvae or pupae at 37°C for two out of every twelve hours over a period of at least two days. Adult females were dissected at least two days after the last heat shock. Flies in which the Gal4-UAS system was used were kept at 29° for 48 hours before dissection unless otherwise noted. The following background stocks were used to generate mitotic clones, which were induced by heat shock at 37° for multiple periods of two hours: RFP-nls, hsflp, FRT19A and hsflp;;FRT82B, RFP-nls / TM3,Sb. Mosaic Analysis with a Repressible Cell Marker (after the method of Lee and Luo) was carried out using GFP-mCD8 (under control of an actin promoter) as the marker.

### Reagents

A list of reagents used in this study is found in Table 2.

**Table 2.** Reagents.

| Reagent | Citation | Vendor or Lab of Origin |
| --- | --- | --- |
| mouse FITC-conjugated anti-α-tubulin | | Sigma Clone DM1A, Lot#114M4817V |
| mouse Alexa fluor 647-conjugated anti-α-tubulin | | Sigma clone DM1A, Lot# 3852427 |
| rabbit anti-Centrosomin | (Lucas and Raff, 2007) | Gift from Raff lab |
| rabbit anti-Inscuteable | (Kraut et al., 1996) | Gift from Knoblich lab |
| Rabbit anti aPKC antibody |  | Santa Cruz Biotechnology Lot#:I2012 |
| Conjugated secondary antibodies |  | ThermoFisher Scientific |
| Fluorescein Phalloidin |  | Invitrogen Lot#:2138394 |
| Alexa fluor 633 Phalloidin |  | Invitrogen Lot#:2274768 |
| Vectashield with DAPI |  | Vector Labs: Lot#:ZH1021 |

### Immunostaining

Ovaries were fixed for 15 minutes in 10% Formaldehyde and 0.2% Tween in Phosphate Buffered Saline (PBS-T), were then incubated in blocking solution (10% Bovine Serum Albumin in PBS) for about one hour at room temperature. Primary and secondary immunostainings lasted at least 3 hours in PBS-T. Three washes of about 10 minutes each in PBS-T were carried out between stainings and after the secondary staining. Primary and secondary antibodies were used at a concentration of 1:150.

### Imaging

Microscopy was performed using either a Leica SP5 point scanning confocal (63x/1.4 HCX PL Apo CS oil lens) or an Andor Dragonfly Spinning Disk confocal microscope (60x water objective). Images were collected with LAS AF or the Andor Fusion software respectively. Minor processing (Gaussian blur) was performed using FIJI.

### Localization Quantification

We used FIJI to perform our quantifications. Starting at the center of the basolateral region (which we defined as 0), a freehand line was drawn around the perimeter of the cell and pixel intensity determined at each point along the line. Background signal, measured in the cytoplasm, was subtracted from these points. Intensities were normalized by dividing the median intensity and positions were normalized to a line with a length of 360. The graphical representations show data points from multiple cells (as indicated in the legend) using a floating 10-point average. Measurement of Mud intensity was sometimes complicated by the proximity between spindle poles, which are highly Mud-positive, and the cortex, which is less so. In these instances, the overlapping region was excluded from the measurement. These exclusions help to explain why Mud intensity profiles are not as smooth as those measured for other proteins.

### Spindle Orientation Measurements

Spindle angle determination was performed as previously described (Finegan et al., 2018). Angles were determined by drawing a first line connecting either the two spindle poles (if applicable) or along the spindle and a second line along the apical surface of the tissue, then measuring the angle between them.

### Statistical analyses

The Mann-Whitney test was used to determine significance when comparing spindle orientation across conditions/genotypes. No statistical method was used to predetermine sample size, the experiments were not randomized, and the investigators were not blinded to allocation during experiments and outcome assessment. The Chi-square test of independence was used to determine significance in Figure 6G.

## References

Berger, J., K.-A. Senti, G. Senti, T.P. Newsome, B. Asling, B.J. Dickson, and T. Suzuki. 2008. Systematic identification of genes that regulate neuronal wiring in the Drosophila visual system. PLoS Genet. 4:e1000085. doi:10.1371/journal.pgen.1000085.

Bergstralh, D.T., H.E. Lovegrove, and D. St Johnston. 2013a. Discs Large Links Spindle Orientation to Apical-Basal Polarity in Drosophila Epithelia. Current Biology. 23:1707–1712. doi:10.1016/j.cub.2013.07.017.

Bergstralh, D.T., H.E. Lovegrove, and D. St Johnston. 2015. Lateral adhesion drives reintegration of misplaced cells into epithelial monolayers. Nat Cell Biol. 17:1497–1503. doi:10.1038/ncb3248.

Bergstralh, D.T., H.E. Lovegrove, I. Kujawiak, N.S. Dawney, J. Zhu, S. Cooper, R. Zhang, and D. St Johnston. 2016. Pins is not required for spindle orientation in the Drosophila wing disc. Development. 143:2573–2581. doi:10.1242/dev.135475.

Bergstralh, D.T., N.S. Dawney, and D. St Johnston. 2017. Spindle orientation: a question of complex positioning. Development. 144:1137–1145. doi:10.1242/dev.140764.

Bergstralh, D.T., T. Haack, and D. St Johnston. 2013b. Epithelial polarity and spindle orientation: intersecting pathways. Philos. Trans. R. Soc. Lond., B, Biol. Sci. 368:20130291. doi:10.1098/rstb.2013.0291.

Bosveld, F., O. Markova, B. Guirao, C. Martin, Z. Wang, A. Pierre, M. Balakireva, I. Gaugue, A. Ainslie, N. Christophorou, D.K. Lubensky, N. Minc, and Y. Bellaïche. 2016. Epithelial tricellular junctions act as interphase cell shape sensors to orient mitosis. Nature. 530:495–498. doi:10.1038/nature16970.

Bowman, S.K., R.A. Neumüller, M. Novatchkova, Q. Du, and J.A. Knoblich. 2006. The Drosophila NuMA Homolog Mud regulates spindle orientation in asymmetric cell division. Dev Cell. 10:731–742. doi:10.1016/j.devcel.2006.05.005.

Brand, A.H., and N. Perrimon. 1993. Targeted gene expression as a means of altering cell fates and generating dominant phenotypes. Development. 118:401–415.

Carminati, M., S. Gallini, L. Pirovano, A. Alfieri, S. Bisi, and M. Mapelli. 2016. Concomitant binding of Afadin to LGN and F-actin directs planar spindle orientation. Nat Struct Mol Biol. 23:155–163. doi:10.1038/nsmb.3152.

Chanet, S., R. Sharan, Z. Khan, and A.C. Martin. 2017. Myosin 2-Induced Mitotic Rounding Enables Columnar Epithelial Cells to Interpret Cortical Spindle Positioning Cues. Curr Biol. 27:3350–3358.e3. doi:10.1016/j.cub.2017.09.039.

Chugh, P., and E.K. Paluch. 2018. The actin cortex at a glance. J Cell Sci. 131. doi:10.1242/jcs.186254.

Culurgioni, S., A. Alfieri, V. Pendolino, F. Laddomada, and M. Mapelli. 2011. Inscuteable and NuMA proteins bind competitively to Leu-Gly-Asn repeat-enriched protein (LGN) during asymmetric cell divisions. 108:20998–21003. doi:10.1073/pnas.1113077108.

Culurgioni, S., S. Mari, P. Bonetti, S. Gallini, G. Bonetto, M. Brennich, A. Round, F. Nicassio, and M. Mapelli. 2018. Insc:LGN tetramers promote asymmetric divisions of mammary stem cells. Nature Communications. 9:1025. doi:10.1038/s41467-018-03343-4.

David, N.B., C.A. Martin, M. Segalen, F. Rosenfeld, F. Schweisguth, and Y. Bellaïche. 2005. Drosophila Ric-8 regulates Galphai cortical localization to promote Galphai-dependent planar orientation of the mitotic spindle during asymmetric cell division. Nat Cell Biol. 7:1083–1090. doi:10.1038/ncb1319.

di Pietro, F., A. Echard, and X. Morin. 2016. Regulation of mitotic spindle orientation: an integrated view. EMBO Rep. e201642292. doi:10.15252/embr.201642292.

Dimitracopoulos, A., P. Srivastava, A. Chaigne, Z. Win, R. Schlomovitz, O.M. Lancaster, M. Le Berre, K. Franze, G. Salbreux, and B. Baum. 2020. Mechanochemical Crosstalk Produces Cell-Intrinsic Patterning of the Cortex to Orient the Mitotic Spindle. Current Biology. 30:3687–3696.e4. doi:10.1016/j.cub.2020.06.098.

Egger, B., J.Q. Boone, N.R. Stevens, A.H. Brand, and C.Q. Doe. 2007. Regulation of spindle orientation and neural stem cell fate in the Drosophila optic lobe. Neural Dev. 2:1. doi:10.1186/1749-8104-2-1.

Finegan, T.M., D. Na, C. Cammarota, A.V. Skeeters, T.J. Nádasi, N.S. Dawney, A.G. Fletcher, P.W. Oakes, and D.T. Bergstralh. 2018. Tissue tension and not interphase cell shape determines cell division orientation in the Drosophila follicular epithelium. EMBO J. e100072. doi:10.15252/embj.2018100072.

Gloerich, M., J.M. Bianchini, K.A. Siemers, D.J. Cohen, and W.J. Nelson. 2017. Cell division orientation is coupled to cell-cell adhesion by the E-cadherin/LGN complex. Nature Communications. 8:13996. doi:10.1038/ncomms13996.

Golub, O., B. Wee, R.A. Newman, N.M. Paterson, and K.E. Prehoda. 2017. Activation of Discs large by aPKC aligns the mitotic spindle to the polarity axis during asymmetric cell division. Elife. 6:166. doi:10.7554/eLife.32137.

Hao, Y., Q. Du, X. Chen, Z. Zheng, J.L. Balsbaugh, S. Maitra, J. Shabanowitz, D.F. Hunt, and I.G. Macara. 2010. Par3 controls epithelial spindle orientation by aPKC-mediated phosphorylation of apical Pins. Curr Biol. 20:1809–1818. doi:10.1016/j.cub.2010.09.032.

Izumi, Y., N. Ohta, K. Hisata, T. Raabe, and F. Matsuzaki. 2006. Drosophila Pins-binding protein Mud regulates spindle-polarity coupling and centrosome organization. Nat Cell Biol. 8:586–593. doi:10.1038/ncb1409.

Johnston, C.A., C.Q. Doe, and K.E. Prehoda. 2012. Structure of an enzyme-derived phosphoprotein recognition domain. PLoS ONE. 7:e36014. doi:10.1371/journal.pone.0036014.

Johnston, C.A., K. Hirono, K.E. Prehoda, and C.Q. Doe. 2009. Identification of an Aurora-A/PinsLINKER/Dlg spindle orientation pathway using induced cell polarity in S2 cells. Cell. 138:1150–1163. doi:10.1016/j.cell.2009.07.041.

Kiyomitsu, T., and I.M. Cheeseman. 2013. Cortical Dynein and Asymmetric Membrane Elongation Coordinately Position the Spindle in Anaphase. Cell. 154:391–402. doi:10.1016/j.cell.2013.06.010.

Kotak, S., C. Busso, and P. Gönczy. 2013. NuMA phosphorylation by CDK1 couples mitotic progression with cortical dynein function. EMBO J. 32:2517–2529. doi:10.1038/emboj.2013.172.

Kraut, R., W. Chia, L.Y. Jan, Y.N. Jan, and J.A. Knoblich. 1996. Role of inscuteable in orienting asymmetric cell divisions in Drosophila. Nature. 383:50–55. doi:10.1038/383050a0.

Liao, E.H., L. Gray, K. Tsurudome, W. El-Mounzer, F. Elazzouzi, C. Baim, S. Farzin, M.R. Calderon, G. Kauwe, and A.P. Haghighi. 2018. Kinesin Khc-73/KIF13B modulates retrograde BMP signaling by influencing endosomal dynamics at the Drosophila neuromuscular junction. PLoS Genet. 14:e1007184. doi:10.1371/journal.pgen.1007184.

Lough, K.J., K.M. Byrd, C.P. Descovich, D.C. Spitzer, A.J. Bergman, G.M. Beaudoin, L.F. Reichardt, and S.E. Williams. 2019. Telophase correction refines division orientation in stratified epithelia. Elife. 8. doi:10.7554/eLife.49249.

Lu, M.S., and K.E. Prehoda. 2013. A NudE/14-3-3 pathway coordinates dynein and the kinesin Khc73 to position the mitotic spindle. Dev Cell. 26:369–380. doi:10.1016/j.devcel.2013.07.021.

Morais-de-Sá, E., and C. Sunkel. 2013. Adherens junctions determine the apical position of the midbody during follicular epithelial cell division. EMBO Rep. 14:696–703. doi:10.1038/embor.2013.85.

Morin, X., and Y. Bellaïche. 2011. Mitotic spindle orientation in asymmetric and symmetric cell divisions during animal development. Dev Cell. 21:102–119. doi:10.1016/j.devcel.2011.06.012.

Nakajima, Y.-I., Z.T. Lee, S.A. McKinney, S.K. Swanson, L. Florens, and M.C. Gibson. 2019. Junctional tumor suppressors interact with 14-3-3 proteins to control planar spindle alignment. Journal of Cell Biology. 218:1824–1838. doi:10.1083/jcb.201803116.

Okumura, M., T. Natsume, M.T. Kanemaki, T.K., 2018. Dynein–Dynactin–NuMA clusters generate cortical spindle-pulling forces as a multi-arm ensemble. eLife doi.org/10.7554/eLife.36559

Olivieri, D., M.M. Sykora, R. Sachidanandam, K. Mechtler, and J. Brennecke. 2010. An in vivo RNAi assay identifies major genetic and cellular requirements for primary piRNA biogenesis in Drosophila. EMBO J. 29:3301–3317. doi:10.1038/emboj.2010.212.

Saadaoui, M., M. Machicoane, F. di Pietro, F. Etoc, A. Echard, and X. Morin. 2014. Dlg1 controls planar spindle orientation in the neuroepithelium through direct interaction with LGN. J Cell Biol. 206:707–717. doi:10.1083/jcb.201405060.

Schaefer, M., A. Shevchenko, and J.A. Knoblich. 2000. A protein complex containing Inscuteable and the Galpha-binding protein Pins orients asymmetric cell divisions in Drosophila. Curr Biol. 10:353–362.

Schiller, E.A., and D.T. Bergstralh. 2021. Interaction between Discs large and Pins/LGN/GPSM2: a comparison across species. Biol Open. 10. doi:10.1242/bio.058982.

Segalen, M., C.A. Johnston, C.A. Martin, J.G. Dumortier, K.E. Prehoda, N.B. David, C.Q. Doe, and Y. Bellaïche. 2010. The Fz-Dsh planar cell polarity pathway induces oriented cell division via Mud/NuMA in Drosophila and zebrafish. Dev Cell. 19:740–752. doi:10.1016/j.devcel.2010.10.004.

Seldin, L., N.D. Poulson, H.P. Foote, and T. Lechler. 2013. NuMA localization, stability, and function in spindle orientation involve 4.1 and Cdk1 interactions. Mol Biol Cell. 24:3651–3662. doi:10.1091/mbc.E13-05-0277.

Siegrist, S.E., and C.Q. Doe. 2005. Microtubule-induced Pins/Galphai cortical polarity in Drosophila neuroblasts. Cell. 123:1323–1335. doi:10.1016/j.cell.2005.09.043.

Siller, K.H., and C.Q. Doe. 2009. Spindle orientation during asymmetric cell division. Nat Cell Biol. 11:365–374. doi:10.1038/ncb0409-365.

Siller, K.H., C. Cabernard, and C.Q. Doe. 2006. The NuMA-related Mud protein binds Pins and regulates spindle orientation in Drosophila neuroblasts. Nat Cell Biol. 8:594–600. doi:10.1038/ncb1412.

Werts, A. 2011. ScienceDirect - Seminars in Cell & Developmental Biology : How signaling between cells can orient a mitotic spindle. Semin Cell Dev Biol.

Wodarz, A., A. Ramrath, A. Grimm, and E. Knust. 2000. Drosophila atypical protein kinase C associates with Bazooka and controls polarity of epithelia and neuroblasts. Journal of Cell Biology. 150:1361–1374.

Wodarz, A., A. Ramrath, U. Kuchinke, and E. Knust. 1999. Bazooka provides an apical cue for Inscuteable localization in Drosophila neuroblasts. Nature. 402:544–547. doi:10.1038/990128.

Yu, F., X. Morin, Y. Cai, X. Yang, and W. Chia. 2000. Analysis of partner of inscuteable, a novel player of Drosophila asymmetric divisions, reveals two distinct steps in inscuteable apical localization. Cell. 100:399–409.

Yu, J.X., Z. Guan, and H.A. Nash. 2006. The mushroom body defect gene product is an essential component of the meiosis II spindle apparatus in Drosophila oocytes. Genetics. 173:243–253. doi:10.1534/genetics.105.051557.

Zajac, A.L., and S. Horne-Badovinac. 2022. Kinesin-directed secretion of basement membrane proteins to a subdomain of the basolateral surface in Drosophila epithelial cells. Current Biology. 32:735–748.e10. doi:10.1016/j.cub.2021.12.025.

Zhao, T., O.S. Graham, A. Raposo, and D. St Johnston. 2012. Growing Microtubules Push the Oocyte Nucleus to Polarize the Drosophila Dorsal-Ventral Axis. Science. 336:999–1003. doi:10.1126/science.1219147.

Zheng, Z., Q. Wan, G. Meixiong, and Q. Du. 2014. Cell cycle-regulated membrane binding of NuMA contributes to efficient anaphase chromosome separation. Mol Biol Cell. 25:606–619. doi:10.1091/mbc.E13-08-0474.

Zhu, J., Y. Shang, C. Xia, W. Wang, W. Wen, and M. Zhang. 2011. Guanylate kinase domains of the MAGUK family scaffold proteins as specific phospho-protein-binding modules. EMBO J. 30:4986–4997. doi:10.1038/emboj.2011.428.

## Drosophila* References (related to Table 1)

Berger, J., Senti, K.-A., Senti, G., Newsome, T. P., Asling, B., Dickson, B. J., & Suzuki, T. (2008). Systematic identification of genes that regulate neuronal wiring in the Drosophila visual system. PLoS Genetics, 4(5), e1000085. http://doi.org/10.1371/journal.pgen.1000085

Bosveld, F., Markova, O., Guirao, B., Martin, C., Wang, Z., Pierre, A., et al. (2016). Epithelial tricellular junctions act as interphase cell shape sensors to orient mitosis. Nature, 530(7591), 495–498. http://doi.org/10.1038/nature16970

Chanet, S., Sharan, R., Khan, Z., & Martin, A. C. (2017). Myosin 2-Induced Mitotic Rounding Enables Columnar Epithelial Cells to Interpret Cortical Spindle Positioning Cues. Current Biology : CB, 27(21), 3350–3358.e3. http://doi.org/10.1016/j.cub.2017.09.039

David, N. B., Martin, C. A., Segalen, M., Rosenfeld, F., Schweisguth, F., & Bellaïche, Y. (2005). Drosophila Ric-8 regulates Galphai cortical localization to promote Galphai-dependent planar orientation of the mitotic spindle during asymmetric cell division. Nature Cell Biology, 7(11), 1083–1090. http://doi.org/10.1038/ncb1319

Kraut, R., Chia, W., Jan, L. Y., Jan, Y. N., & Knoblich, J. A. (1996). Role of inscuteable in orienting asymmetric cell divisions in Drosophila. Nature, 383(6595), 50–55. http://doi.org/10.1038/383050a0

Liao, E. H., Gray, L., Tsurudome, K., El-Mounzer, W., Elazzouzi, F., Baim, C., et al. (2018). Kinesin Khc-73/KIF13B modulates retrograde BMP signaling by influencing endosomal dynamics at the Drosophila neuromuscular junction. PLoS Genetics, 14(1), e1007184. http://doi.org/10.1371/journal.pgen.1007184

Lovegrove, H. E., Bergstralh, D. T., & St Johnston, D. (2019). The role of integrins in Drosophila egg chamber morphogenesis. Development, 146(23). http://doi.org/10.1242/dev.182774

Perkins, L. A., Holderbaum, L., Tao, R., Hu, Y., & Sopko, R. (2015). The transgenic RNAi project at Harvard Medical School: resources and validation. Genetics, 201(3), 843–852. http://doi.org/10.1534/genetics.115.180208

Perrimon, N. (1988). The maternal effect of lethal(1)discs-large-1: a recessive oncogene of Drosophila melanogaster. Developmental Biology, 127(2), 392–407. http://doi.org/10.1016/0012-1606(88)90326-0

Woods, D. F., & Bryant, P. J. (1989). Molecular cloning of the lethal(1)discs large-1 oncogene of Drosophila. Developmental Biology, 134(1), 222–235.

Yu, F., Morin, X., Cai, Y., Yang, X., & Chia, W. (2000). Analysis of partner of inscuteable, a novel player of Drosophila asymmetric divisions, reveals two distinct steps in inscuteable apical localization. Cell, 100(4), 399–409.

Yu, J. X., Guan, Z., & Nash, H. A. (2006). The mushroom body defect gene product is an essential component of the meiosis II spindle apparatus in Drosophila oocytes. Genetics, 173(1), 243–253. http://doi.org/10.1534/genetics.105.051557

Zajac, A. L., & Horne-Badovinac, S. (2022). Kinesin-directed secretion of basement membrane proteins to a subdomain of the basolateral surface in Drosophila epithelial cells. Current Biology, 32(4), 735–748.e10. http://doi.org/10.1016/j.cub.2021.12.025

## Reagent References – Related to Table 2

Lucas, E.P., and J.W. Raff. 2007. Maintaining the proper connection between the centrioles and the pericentriolar matrix requires Drosophila centrosomin. Journal of Cell Biology. 178:725–732. doi:10.1083/jcb.200704081.

